# Nanocomposite bioink exploits dynamic covalent bonds between nanoparticles and polysaccharides for precision bioprinting

**DOI:** 10.1101/839985

**Authors:** Mihyun Lee, Kraun Bae, Clara Levinson, Marcy Zenobi-Wong

## Abstract

The field of bioprinting has made significant recent progress towards engineering tissues with increasing complexity and functionality. It remains challenging, however, to develop bioinks with optimal biocompatibility and good printing fidelity. Here, we demonstrate enhanced printability of a polymer-based bioink based on dynamic covalent linkages between nanoparticles (NPs) and polymers, which retains good biocompatibility. Amine-presenting silica NPs (ca. 45 nm) were added to a polymeric ink containing oxidized alginate (OxA). The formation of reversible imine bonds between amines on the NPs and aldehydes of OxA lead to significantly improved rheological properties and high printing fidelity. In particular, the yield stress increased with increasing amounts of NPs (14.5 Pa without NPs, 79 Pa with 2 wt% NPs). In addition, the presence of dynamic covalent linkages in the gel provided improved mechanical stability over 7 days compared to ionically crosslinked gels. The nanocomposite ink retained high printability and mechanical strength, resulting in generation of centimetre-scale porous constructs and an ear structure with overhangs and high structural fidelity. Furthermore, the nanocomposite ink supported both *in vitro* and *in vivo* maturation of bioprinted gels containing chondrocytes. This approach based on simple oxidation can be applied to any polysaccharide, thus the widely applicability of the method is expected to advance the field towards the goal of precision bioprinting.

## 1. Introduction

Bioprinting has shown great promise and potential as a versatile technology to generate 3D tissue constructs closely resembling the multiscale and multicellular architecture as well as mechanical heterogeneity of native tissue.^[1]^ Such bioprinted tissues can provide patient-tailored grafts, *in vitro* drug resistance testing systems as well as high throughput drug screening platforms with high precision and reproducibility.^[1, 2]^ Fabrication of cell-laden constructs is carried out either by direct printing of cells with/without supporting materials or seeding cells on a printed scaffold. The former approach is advantageous to achieve high cell density, continuous distribution of cells similar to native tissue, and precise positioning of different cell types. An ink containing living cells is called ‘bioink’, which is distinct from biomaterials inks consisting of polymers or precursors of hydrogels containing biological factors.^[3]^ Development of a bioink with both high biocompatibility and printability has been challenging because only limited materials, crosslinking methods and processing conditions permit high cell viability and function after bioprinting. Most traditional hydrogels have shown good biocompatibility but suffer from low printability, printing fidelity and/or mechanical stability after printing ^[4]^.

In extrusion-based bioprinting, printability of a bioink mainly depends on its rheological properties such as shear-thinning and shear-recovery properties and the presence of a yield stress. High shear-thinning properties enable an ink with high viscosity to flow through a needle, and a high shear-recovery rate and yield stress allow the ink to keep hold and recover its shape after the extrusion phase. A great deal of effort have been made to modulate the rheological properties of hydrogel-based bioinks along these lines.^[5]^ Addition of nano/microparticles is one of the most widely adopted methods to improve rheological properties of a polymer-based hydrogel. ^[6, 7]^ In particular, nanofibrils or NPs retaining intrinsic shear-thinning properties such as nanocellulose and 2D silicates have been utilized to prepare bioinks in combination with various polymers including gelatin,^[8]^ poly(ethylene glycol)(PEG),^[9]^ carrageenan,^[10]^ hyaluronic acid (HA),^[11]^ alginate,^[12]^ and fibrin.^[13]^ The mechanical properties and printability of such nanocomposite bioinks mainly depend on the properties of the NP component in the ink, which have shown only limited mechanical strength/stability and printing quality to date. ^[8–13]^ Furthermore, those shear-thinning NPs are biologically inactive or retain limited biological activities towards a specific type of tissue. For example, 2D silicates have been intensely utilized for printing orthopedic tissue due to its known osteogenic activity ^[14]^. However, there is no available shear-thinning NPs which retain bioactivities towards other types of tissue than orthopedic ones. Considering a wide variety of available therapeutic NPs such as drug-encapsulating liposomes or iron NPs, ^[15, 16]^ diversification of the nano/microparticles in nanocomposite-based bioinks would facilitate the production of various types of bioprinted tissue analog retaining desirable biological functions.

In the field of injectable and/or self-healing nanocomposite hydrogels, there have been great advances to formulate functional nanocomposite hydrogels where interactions between polymers and NPs were actively modulated to achieve viscoelastic properties not achievable by polymers or NPs alone.^[17–19]^ For example, addition of β-cyclodextrin surface-modified silica NPs (SiNPs) to adamantine-functionalized PEG resulted in greatly increased viscosity at a low shear rate (157 Pa s for the native hydrogel and ca.1300 Pa s for the hydrogel with the NPs) as well as improved shear-thinning properties via host-guest interactions between β-cyclodextrins and adamantines.^[20]^ In another example, a self-healing hydrogel was prepared by mixing dendritic macromolecules (G*n*-binders) and nanoclays modified with sodium polyacrylate.^[21]^ The resultant hydrogel exhibited moldability as well as exceptionally high mechanical strength, which was attributed to reversible interactions between the oxyanions on the nanoclays and guanidinium ions on the G*n*-binders. Despite such promising advances in controlling viscoelastic and mechanical properties of nanocomposites, the strategies to date have rarely been adopted for formulating bioinks.

We recently reported that incorporation of solid SiNPs into a polymer-based bioink could lead to significant enhancement of rheological properties and printing fidelity.^[22]^ The enhancement was mainly due to electrostatic interactions between cationic SiNPs and anionic polysaccharides, and was greatly affected by the size and concentration of NPs as well as the length of polymer chains. As a result a centimeter-scale ear construct was printed with high printing fidelity as well as good biocompatibility.

Here, a novel approach for preparing nanocomposite bioinks is reported where formation of dynamic covalent linkages between SiNPs and oxidized polysaccharides provides desirable rheological properties for high-precision bioprinting without compromising *in vitro* and *in vivo* biocompatibility. In the present study, the reversible imine bond is chosen as a dynamic covalent linkage because it has been extensively utilized in various biocompatible hydrogels as bond formation can take place at physiological pH with no additional stimuli.^[23]^ In addition, aldehyde-presenting polymers can be easily prepared via oxidation of polysaccharides such as alginate,^[24]^ dextran,^[25]^ HA,^[26]^ gellan ^[27]^ and cellulose^[28]^. To demonstrate the concept, aminopropyl-modified SiNPs (NH_2_-SiNPs) were added to a mixture of polymers containing oxidized alginate (OxAlg), (Figure 1a). Addition of the NH_2_-SiNPs resulted in enhanced shear thinning properties and increased yield stress compared to the polymer mixture without the NPs, which lead to improved printability as well as high structural fidelity. Compared to electrostatic interactions^[22]^ used in our previous study where 6 wt% cationic SiNPs were added to an anionic polymer ink to obtain desirable rheological enhancement for printing, the dynamic covalent linkages led to significant improvement in rheological properties as well as printability at a lower concentration of NPs (2 wt%). In addition, the bioprinted gels proceeded *in vitro* and *in vivo* maturation.

**Figure 1.**
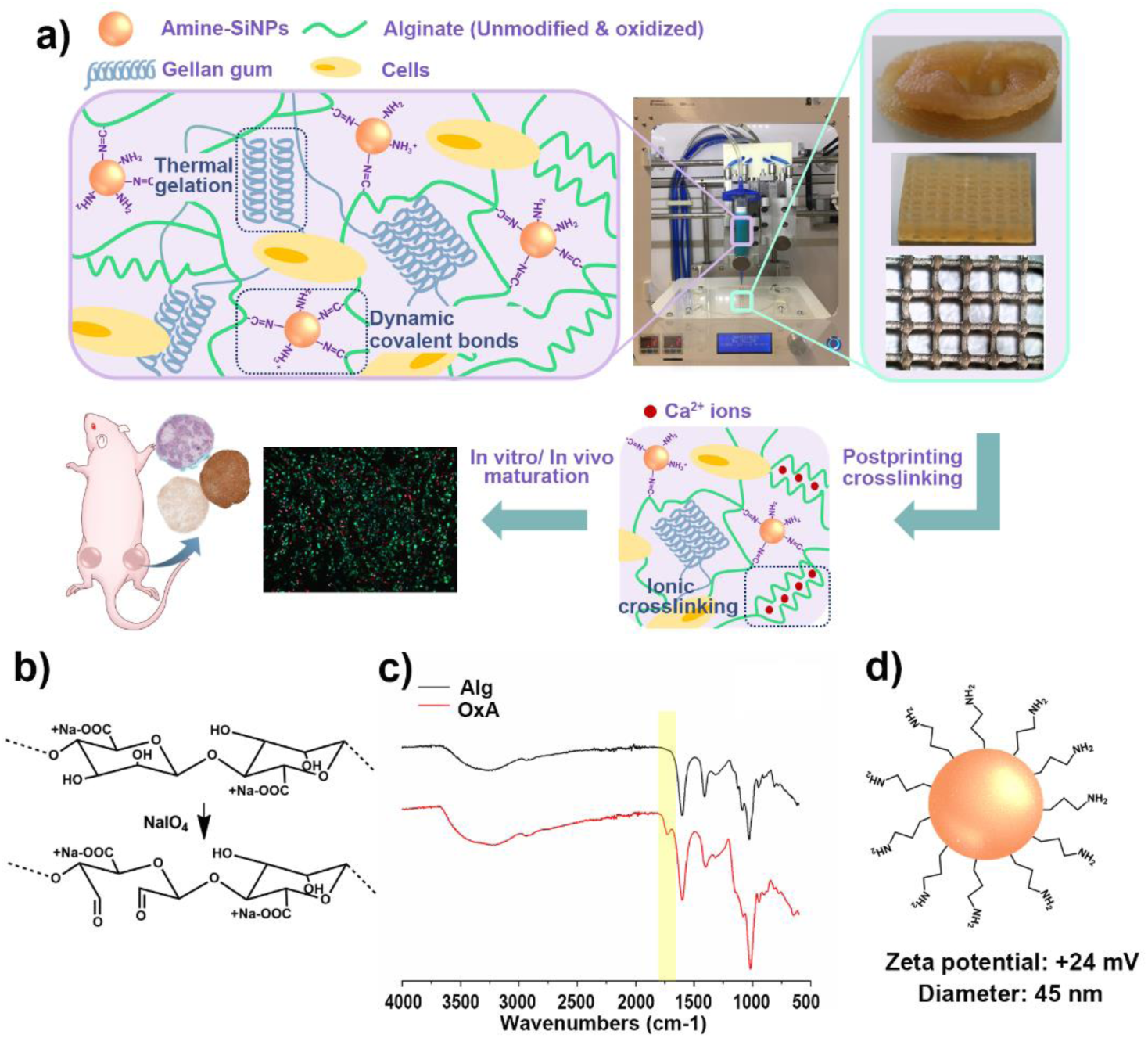
(a) Schematic presentation of the nanocomposite bioink based on dynamic covalent imine bonds between SiNPs and oxidized polysaccharides and bioprinting with the bioink. (b) Synthesis of oxidized alginate using sodium periodate as an oxidizing reagent. (c) FTIR spectra of unmodified (Alg) and oxidized (OxA) alginate. (d) Schematic presentation of a surface modified SiNP used in this study.

## 2. Materials and Methods

### 2.1 Aminopropyl-modification of SiNPs

Aminopropyl-modification of SiNPs was performed according to previously reported methods with modifications. [29, 30] The SiNP dispersion (LUDOX^®^ TM-50 colloidal silica, Sigma-Aldrich, Buchs, Switzerland) was diluted with ultrapure water (30 mL LUDOX^®^ TM50 to 400 mL water) followed by addition of ammonia (10 mL) (NH3, 28-30%, Merck, Darmstadt, Germany) and (3-aminopropyl)trimethoxysilane (24 mL) (APTMS 97%, Sigma-Aldrich). The mixture was incubated in an oil bath at 40 °C with magnetic stirring at 400 rpm for 16 h. Afterwards, the excess reagents were removed by repeated centrifugation at 15,000 G for 20 min. During the washing step, re-dispersion of the pellet was possible by adjusting the pH to ~1 and induction of pellet formation was done at ~pH 10. The NPs were washed three times in ethanol (99 % absolute, Merck) and three times in deionized water. The modified SiNP dispersion was kept at pH 2-3 with gentle magnetic stirring at room temperature. The size and zeta potential were analyzed using a zetasizer (Zetasizer Nano ZS, Malvern Instrument Ltd, UK).

### 2.2 Preparation of OxAlg and depolymerized alginate

Sodium alginate (I3-G, Kimica corporation, Tokyo, Japan) was dialyzed against dionized water for >72 h for purification followed by lyophilization. OxAlg was synthesized according to the method reported by Wang et al ^[24]^. Briefly, 1 g sodium alginate was dissolved in 100 mL ultrapure water with magnetic stirring at 300 rpm overnight. 0.6 molar equivalent amounts of sodium periodate (Sigma-Aldrich) was added to the sodium alginate solution. The mixture was magnetically stirred in the dark at room temperature for 5 h, followed by addition of 3 mL ethylene glycol (Sigma-Aldrich) to remove unreacted sodium periodate. After another hour of magnetic stirring, the mixture was dialyzed against ultrapure water for 72 h using dialysis tubing (MWCO 12 kDa, Sigma-Aldrich) followed by lyophilization. The degree of oxidation was determined by iodometry. ^[24]^ A solution of 10 % sodium bicarbonate (10 mL)(Sigma-Aldrich) and 20 % potassium iodide (2 m L)(Sigma-Aldrich) was prepared, to which 5 mL of the reaction mixture was added. The solution was stirred in dark for 30 min. The liberated iodine was quantified via titration with 0.1 M sodium thiosulfate solutuion (Sigma-Aldrich) using a 1 wt% starch (Sigma-Aldrich) solution as an indicator. The molarity of unreacted sodium periodate was calculated from the detected amounts of thiosulfate, based on which the degree of oxidation was calculated.

To prove that the formation of imine bonds in the NC ink led to rheological enhancement, we prepared control inks which have the same composition to their counterpart (Polym or NC ink) but do not contain aldehydes. Fot this, alginate with a similar molecular weight to OxAlg was prepared through depolymerization of intact alginate and used for the preparation of the control inks instead of OxAlg. Depolymerization of alginate was performed according to a previously reported method. ^[31]^ Briefly, sodium alginate was dissolved in ultrapure water at 2 % (w/v) followed by an adjustment of pH to 4.4 using a 1M HCl solution. The solution was then placed in an oil bath at 80 °C with magnetic stirring at 400 rpm. The depolymerization was stopped after 30 h, when the depolymerized alginate was eluted at the same retention time as OxAlg in gel permeation chromatography (GPC) analyses. (Figure S2) The depolymerized alginate was dialyzed against deionized water for 72 h followed by lyophilization.

### 2.3 Preparation of inks

To prepare the polymeric ink, gellan (350 mg, referred to as 3.5%)(KELCOGEL, Low acetylated gellan gum, CP Kelco, Atlanta, GA, USA), alginate (150 mg, referred to as 1.5 %) and OxAlg (150 mg, referred to as 1.5 %) were added to a buffer solution (10 mL) (4-(2-hydroxyethyl)-1-piperazineethanesulfonic acid (HEPES) 20 mM at pH 7.4 containing glucose 300 mM). The mixture of polymer powder and the buffer solution was magnetically stirred at 200 rpm in an oil bath at 90 °C for 1 h. Afterwards, the solution was cooled down to room temperature with mixing using a spatula for 12 min. For preparation of the nanocomposite ink, everything else was done as for the polymeric ink except NH_2_-SiNP dispersion in the same buffer solution (HEPES 20 mM at pH 7.4 containing glucose 300 mM) was used. To remove any agglomerations, both inks were further homogenized using two syringes connected by a lure connector. A helical mixing component (Medmix Systems AG, Risch-Rotkreuz, Switzerland) was placed in the connector, and the ink was passed between the two syringes through the connector at ~0.5 Hz for 10 min. Afterwards, the inks were centrfuged at 3000 rpm for 90 min to remove air bubbles. The prepared inks were then kept at 4 °C.

### 2.4 Mechanical testing

All rheological characterization was done using the Anton Paar MCR 301 rheometer (Anton Paar, Zofingen, Switzerland) at 25 °C with a gap distance of 0.5 mm. For a shear-thinning test, the shear rate was logarithmically increased in a range of 0.01 – 500 s^−1^ and the viscosity was recorded. For assessing shear recovery properties, storage (G’) and loss (G”) moduli were measured at the frequency of 1 rad/s with 0.5 % strain for 10 min, which was followed by measurement at the same frequency with 500 % strain for 10s. This high strain-low strain transition was repeated two times. For measurement of yield stress, G’ and G” were recorded at varying shear stress ranged from 0.1 to 100 Pa. The yield stress was determined as the crossover point of G’ and G”.

Compressive and tensile moduli were measured using a texture analyzer (Stable Micro Systems, London, England) for gels crosslinked in a solution containing 100 mM CaCl_2_ and 70 mM NaCl for 1 h. For compression tests, gels with 1 mm in thickness and 4 mm in a diameter were used, and for tensile tests, specimen with a dimension of 15 * 2.3 * 1 mm (length *width*thickness) were prepared. Both compression and tensile tests were carried out at a speed of 0.01 mm/s, and the moduli were calculated from slopes in linear-viscoelastic (LVE) ranges (0-4 % strain).

### 2.5 3D printing

3D printing for the assessment of printability was performed using the INKREDIBLE+ 3D printer (Cellink, Gothenburg, Sweden) and printing needles with a diameter of 410 μm and conical shape. 3D objects were designed with Autodesk Fusion 360 software, and g-codes were created using Slic3r software. The printing pressures for the polymeric and nanocomposite inks were ~25 kPa and ~36 kPa, respectively. Typical printing speed was 10 mm/s. Diffusion rates and collapse area factors were calculated according to previous studies by Habib et al^[32]^.

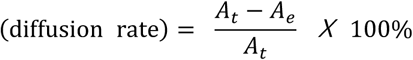

where *A*_*t*_ is the theoretical area of a lattice and *A*_*e*_ is the experimentally measured area of a lattice, and

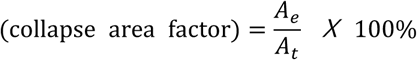

where *A*_*t*_ is the theoretical area delineated between two adjacent pillars (distance x height of pillar) and *A*_*e*_ the area delineated between the printed line connecting the pillars. For printing of cell-laden gels, the Biofactory bioprinter (RegenHu, Switzerland) within a sterile hood was used. The printer was sterilized by UV irradiation for 1 h. For preparation of bioinks, cell suspension (1*10^8^ cells/mL) was mixed into each ink at the ratio of 1:9 (v/v) yielding a final concentration of 1*10^7^ cells/mL. After printing, the objects were crosslinked in a calcium-containing solution (CaCl_2_ 100mM, NaCl 70 mM) for 30 min. The crosslinked gels were washed using a 0.9 % NaCl solution and cultured in the incubator at 37 °C and 5 % CO_2_.

### 2.6 Isolation and culture of chondrocytes

All biological reagents were purchased from Life Technolgies (Zug, Switzerland) unless otherwise stated. Bovine knee cartilage was obtained from a local butcher’s shop (Metzgerei Angst AG, Zurich, Switzerland). Cartilage was collected from condyle and minced into small cubes (ca. 1*1*1 mm), followed by digestion with collagenese (9.6 mg for 1 g of cartilage) at 37 °C for 5 h. After digestion, the digestion mixture was squentially filtered through sieves with mesh sizes of 100 μm and 70 μm. The filtered solution was centrifuged at 250 G for 8 min, followed by washing with once with expansion medium once. The cells were dispersed in recovery media, frozen and stored in liquid nitrogen. For expansion of cells, a high-glucose medium supplemented with 10 % fetal bovine serum, 50 μg/mL ascorbic acid (Sigma-Aldrich), and 10 μg/mL gentamycin was used. For chondrogenesis, cells were cultured in a medium containing 1 % ITS premix (Corning Life Sciences, Amsterdam, The Netherlands), 50 μg/mL ascorbic acid, 25 μg/mL L-proline (Sigma-Aldrich), 10 μg/mL gentamycin and 10 ng/mL TGF-β3 (Peprotech, London, UK). For the use of human chondrocytes, patient consent and ethical approval were obtained (KEK-ZH 2017-02101). The isolation of human chondrocytes was done according to the protocol used for bovine articular chondroytes with slight modification. Briefly, cartilage were treated with pronase (0.5 %) for 1.5 h, and minced into small cubes. The cubes were treated with a 0.1 % collaganese solution overnight followed by sequential filtration with seives. The isolated cells were maintained in a DMEM medium suplemented with 20 % FBS, 50 μg/mL ascorbic acid and antibiotic-antimycotic. Viability was assessed by live/dead staining where cells were incubated in a solution containing calcein-AM (2 μM) and propidium iodide (5.88 μg/mL) followed by washing with phosphate buffered saline (PBS). Imaging was done using a microscope equipped with apotome (Axio Observer, Carl Zeiss, Oberkochen, Germany).

### 2.7 In vivo implantation

After 4 weeks of preculture in chondrogenic media, the bioprinted gels (6 gels per condition, 2 gels per mouse) were implanted subcutaneously in female, 3-month old nude mice (Charles River). The *in vivo* study was performed in compliance with the ethical guidelines of the Veterinary Office of the Canton of Zurich (application number ZH118/2017). Briefly, 5 % isoflurane was used to anesthetize the mice and and Metacam (2 mg/kg) was injected before surgery to prevent post-surgical pain. Two subcutaneous pockets were created laterally to the dorsal midline, by making an incision at the level of the hip joint and clearing the fascia. After 8 weeks *in vivo*, the gels were explanted after euthanizing the mice by CO_2_ asphyxiation.

### 2.8 Histology

Samples were fixed by immersion in 4 % formaldehyde for 1 h at 4°C, dehydrated in baths of gradually increasing ethanol concentration and embedded in paraffin. 5 µm-thick sections were deparaffinized in xylene and rehydrated in baths of decreasing concentration of ethanol prior to staining. Safranin O/Fast green, Alizarin Red and Hematoxylin & Eosin (H&E) staining was performed according to standard protocols. For immunohistochemical staining of collagens, the sections were treated with hyaluronidase (Sigma-Aldrich, 1200U/ml, 30 min at 37°C) for antigen retrieval and with normal goat serum (NGS, Abcam, 5 % in PBS for 1 h at room temperature) for blocking. Primary antibodies, both rabbit anti-human (Rockland 600-401-104, 1:200, for collagen II, Abcam ab138492, 1:1500 for collagen I) were applied in 1 % NGS, overnight at 4 °C. The goat anti-rabbit IgG-HRP (Abcam, ab6721, 1:1500 in 1 % NGS) was added after washing and quenching of the endogenous peroxidase activity (1 % H_2_O_2_ in water for 20 min) at room temperature for 1 h. Antibody detection was done with the DAB kit (Abcam, 4 min incubation with the chromogen) as per manufacturer’s protocol. Slides were dehydrated and mounted (Eukitt mounting medium) prior to imaging with a slide scanner (Pannoramic 250 Flash II).

### 2.9 Statistical analyses

Statistics was performed using OriginPro 8 software. Two sample t-test was performed to calculate p-values. P-values < 0.05 were considered to be significant statistically.

## 3. Results

### 3.1 Preparation and characterization of NH_2_-SiNP and OxAlg

OxAlg was prepared via oxidizing alginate by sodium periodate (Figure 1b). The presence of aldehyde groups after oxidation was confirmed by Attenuated Total Reflectance-Fourier Transform IR Spectroscopy (ATR-FTIR), where the carbonyl stretch at 1720 cm^−1^ appeared after oxidation (Figure 1c). The degree of oxidation was determined by iodometry to be 0.6. Solid SiNPs were chosen because they have rarely been used in bioprinting^[33]^ due to lack of intrinsic shear-thinning properties despite of their high biocompatibility,^[15, 34]^ well-established protocols for controlled synthesis of size, porosity and surface chemistry.^[35, 36]^ The surface of SiNPs was modified with aminopropyl moieties to render the particle surfaces to present primary amines which can form imine linkages with aldehydes. (Figure 1d). The zeta potential and size of the NH_2_-SiNP were +24 mV and 45 nm, respectively.

### 3.2 Preparation and mechanical characterization of bioinks

Nanocomposite inks were modified from a previously reported bioink comprised of alginate and gellan which showed moderate printability/printing fidelity and high biocompatibility.^[37]^ Here, NH_2_-SiNPs and OxAlg were added to the alginate/gellan-based ink to prepare the nanocomposite inks with dynamic covalent linkages (Figure 1a). For comparison, a control ink was also prepared using the same composition to the nanocomposite ink but without NH_2_-SiNPs.

First, the ratio between OxAlg and alginate in a nanocomposite ink was optimized. Keeping the total concentration of OxAlg and alginate constant at 3 %, the ratio of OxAlg to alginate was varied from 3:0 to 1.5:1.5 (%). The concentration of gellan was kept constant (3.5 %) for all conditions. For nanocomposite inks, NH_2_-SiNPs were added at 2 wt% to the control inks. The inks with the ratio of 3/0 (OxAlg/alginate, i.e. 3 % OxAlg and 0 % alginate) exhibited inhomogeneous crosslinking as well as prominent phase separation as shown in Figure S1. It was not possible to extrude a uniform strand from both inks with/without NH_2_-SiNPs due to the presence of over-crosslinked clumps of gel (Figure 2a). Decreasing the ratio of OxAlg to alginate greatly alleviated the phase separation as well as over-crosslinking (Figure 2a). The inks with the ratio of 1.5/1.5 allowed the inks to be extruded without causing any clogging. Therefore, we chose the ratio of OxAlg to alginate 1.5:1.5. The polymeric and nanocomposite inks with this ratio of OxAlg to alginate (1.5:1.5) are referred to as *Polym ink* and *NC ink*.

**Figure 2.**
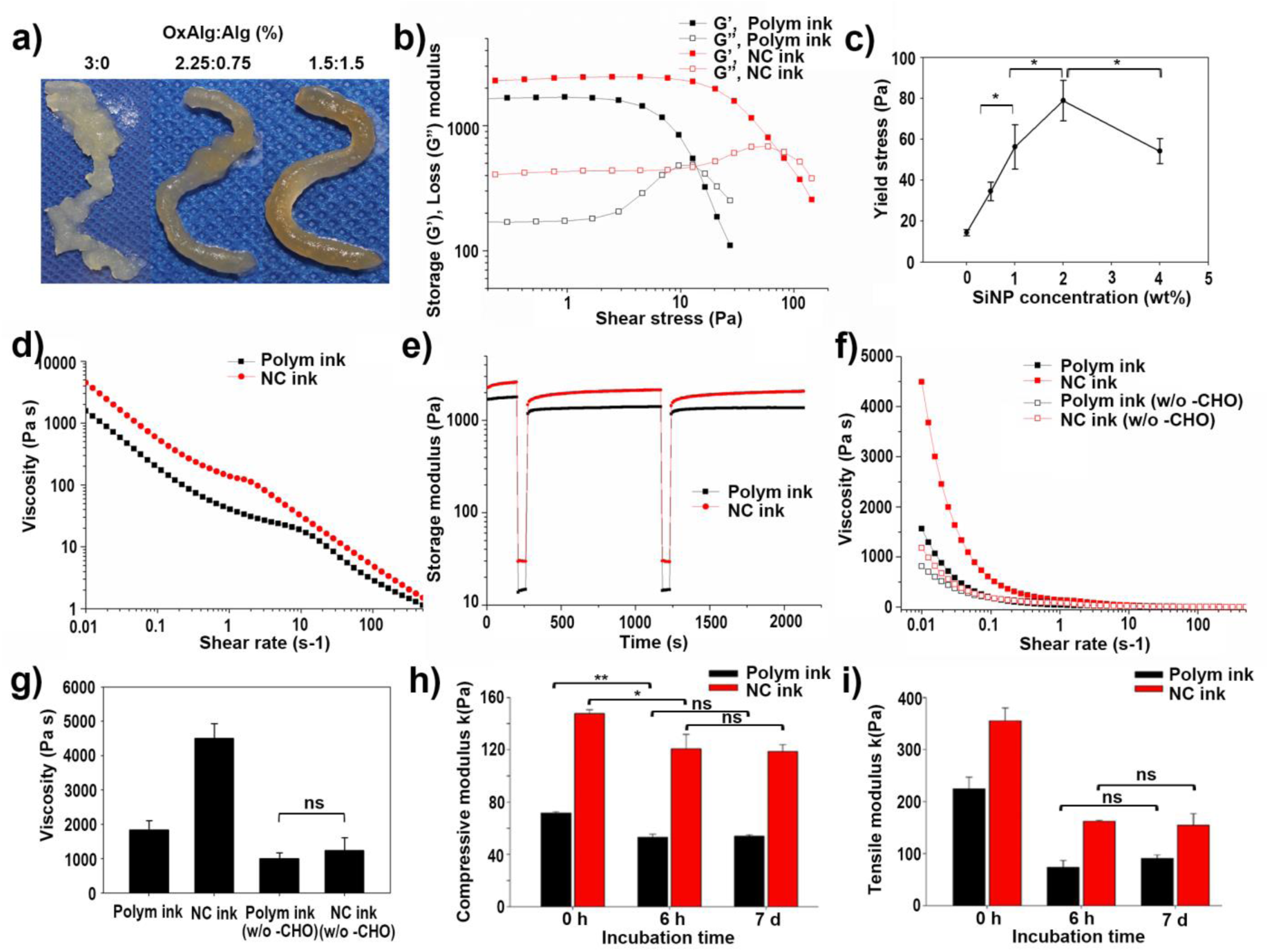
Mechanical properties of the polymeric (Polym) and nanocomposite (NC) inks. (a) Photographs of NC inks with different ratios of OxAlg to Alg. (b) Storage and loss modulus as a function of % strain. (c) Yield stress of NC inks with different amounts of NPs. (d) Shear thinning properties. (e) Shear recovery properties. (f,g) Viscosities at a range of shear rates (f) and at the shear rate of 0.01s^−1^ of the Polym inks (with, without OxA) and NC inks (with, without OxA) (g). (h,i) Compressive (h) and tensile (i) modulus of calcium crosslinked Polym gels and NC gels at different incubation time. * P value <0.05, ** P value <0.01, ns: Not significantly different

Next, rheological properties were characterized. First, addition of NH_2_-SiNPs to the Polym ink led to increased yield stress from 14.5 Pa without NPs to 79 Pa with 2 wt% NH_2_-SiNP (446 % increase)(Figure 2b). Addition of only 0.5 wt% NH_2_-SiNP resulted in 139 % increased yield stress, and the yield stress increased with increasing amounts of NH_2_-SiNP from 0.5 to 2 wt% (Figure 2c). Further increase in the NH_2_-SiNP concentration to 4 wt% did not lead to an additional increase in yield stress. Therefore the NC ink with 2 wt% NH_2_-SiNP was used for the rest of the study.

Shear thinning and recovery properties were characterized. The NC ink retained higher viscosity at the low shear rate (0.01 s-1) compared to the Polym ink and decrease to almost the same viscosity as the Polym ink at the high shear rate (500 s-1), which implies enhanced shear thinning properties. (Figure 2d) In addition, both inks recovered storage modulus after sequential high shear rate phases (Polym Ink 77 % and NC ink 80 % recovery after the second high shear phase). Interestingly, the damping factor (G”/G’) of the Polym ink tended to increase after high shear rate phases (147 % of the initial value after the second high shear rate phase), whereas for the NC ink the same trend was not observed (80 % after the second high shear rate phase) (Figure 2e).

We further studied if the observed rheological enhancement was indeed a result of the formation of imine linkages between NH_2_-SiNPs and OxAlg. For that, Polym and NC inks were prepared using degraded alginate instead of OxAlg which has a similar molecular chain length to OxAlg (Figure S2) without the aldehyde groups. As a result, addition of NH_2_-SiNP to the Polym ink made of degraded alginate without aldehydes did not lead to significant increases in the viscosity or shear thinning properties (Figure 2f,g), which confirms that the presence of both aldehydes and NH_2_-SiNPs were required for rheological enhancement.

Next, mechanical stiffness after ionic crosslinking by calcium ions was studied (Figure 2h,i). The NC gel retained higher compressive (106 % increase) and tensile (58 % increase) moduli compared to the Polym gel. For both Polym and NC gels, compressive and tensile moduli significantly decreased after 6 h. However, the NC gel exhibited an improved mechanical stability at day 7 where the NC gel retained 120 % higher compressive modulus and 70 % higher tensile modulus compared to the Polym gel.

### 3.3 Assessment of printability and printing fidelity

For assessment of printability, different printing pressures were used for the Polym and NC ink, which were the minimum pressures that induced continuous extrusion of strands without breakage for >30 s for each ink. At the fixed printing pressure, stability and thickness of extruded strands were anayzed. (Figure 3a). For both inks, extrusioin at printing speed of 2 mm/s resulted in poorly defined strand structures due to too high feed rates. With increased printing speed at 5 mm/s, printing individual strands with a uniform width was possible. Further increases in printing speed resulted in decreased line widths. (Figure 3b,c) At given printing speed the NC ink resulted in a thinner line width than the Polym ink. For example, at speed 10 mm/s, the line width of the NC and Polym ink were 397 μm and 522 μm, respectively. For both inks, printing at speed higher than 15 mm/s led to occasional breakage of strands, which suggests the range of stable strand extrusion to be ca. 5 – 10 mm/s. Therefore, printing speed of 10 mm/s was used for next steps.

**Figure 3.**
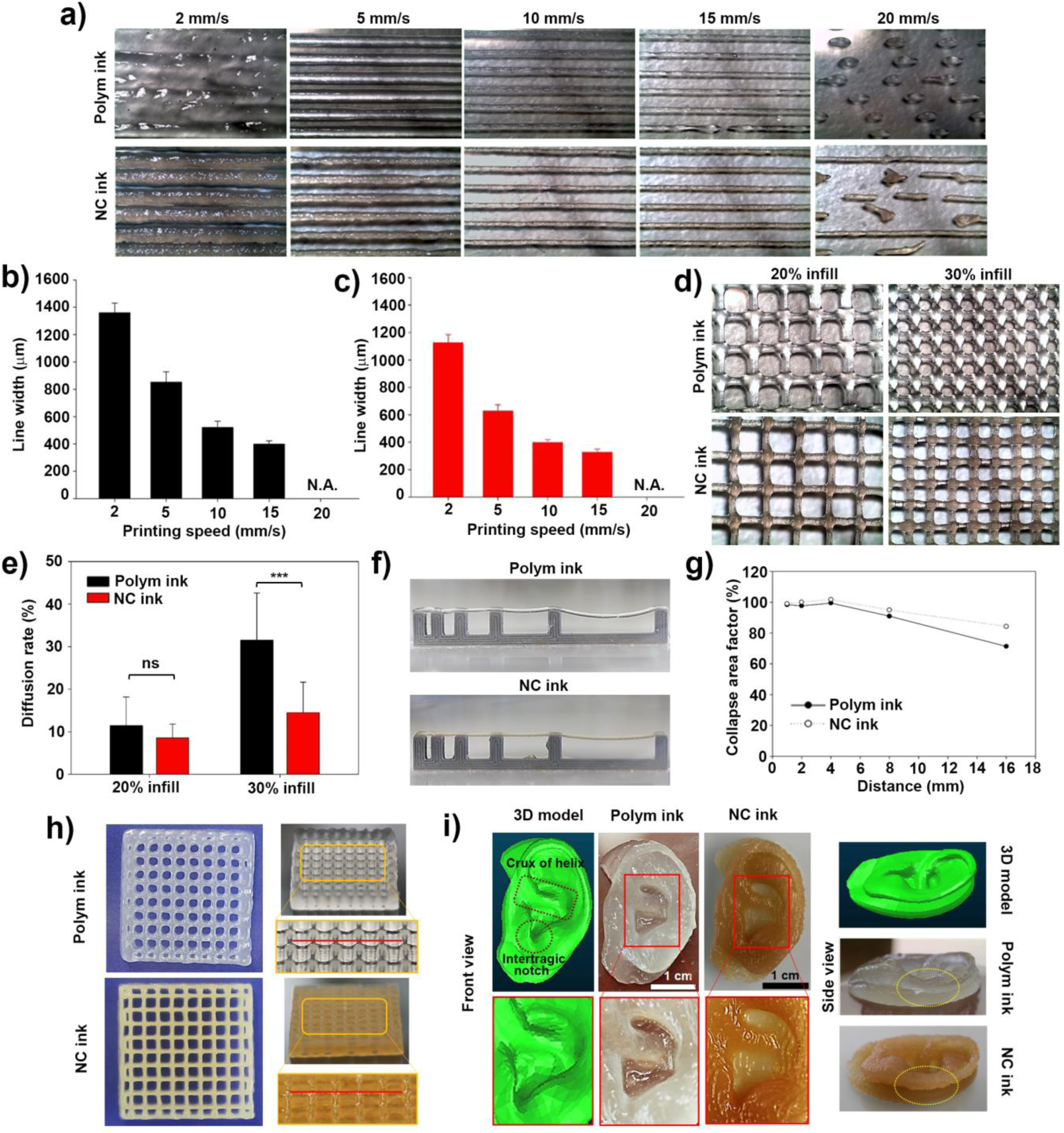
Printability and printing fidelity of the inks. (a) Photographs of printed lines at different printing speed. (b,c) Widths of lines printed using the Polym ink (b) or NC ink (c). (d) Photographs of printed grid structures with different spacing. (e) Diffusion rate of each ink for grid structures with different spading. (f) Photographs of printed strands on a construct with pillars. (g) Collapse area factors of the Polym ink and NC ink as a function of a distance between pillars. (h) Photographs of 10-layerd grid structures. Yellow box indicates sagging of the Polym strand, and good spanning properties of the NC strands. (i) Photographs of ear structures. Red box indicates magnification ***P<0.001

To assess printing fidelity, grid structures with different pore sizes were printed. (Figure 3d) For both inks, when a grid with a large pore size (20 % infill) was printed, the angular pore shape was maintained. However, printing a grid with a small pore size (30 % infill) using the Polym ink resulted in significant diffusion of materials at the intersections, which led to loss of the angular shape of pores. In contrast, the grid structure printed using the NC ink retained the angular shape and showed high printing fidelity. Indeed, the calculated diffusion rate of the NC ink was significantly higher than the value of the Polym ink for the structure with 30 % infill. (Figure 3e)

Additionally, the ability of the strands to span pillars spaced at increasing distances (1, 2, 4, 8, 16 mm) was measured (Figure 3f). Degrees of the collapse were affected by both the ink type and distance between pillars. For small spacing (1 – 4 mm), no notable collapse was observed for both inks. The Polym ink strand printed between pillars 8 mm apart from each other collapsed slightly. For spacing of 16 mm, the strands collapsed significantly, and the calculated collapse area factor was lower for the Polym ink (71 %) than that of the NC ink (84 %)(Figure 3g).

Finally, we printed centimeter-scale structures to study how diffusion of material and collapse affected structual fidelity of a large construct. When a grid structure with 10 layers was printed using the Polym ink, the angular pore shape was mostly lost as observed in the 2-layer grid structure (Figure 3h, top, left), and sagging of printed strands was clearly observed (Figure 3h, top, right). Both ink diffusion and collapses were greatly suppressed when the same structure was printed using the NC ink, which resulted in high structural fidelity (Figure 3h, bottom). In addition, a centimeter-scale ear was printed as a clinically relevant structure. As shown in Figure 3i, the ear printed using the NC ink showed higher structural similarity to the 3D digital model compared to the ear printed using the Polym ink. For instance, the ear printed using the Polym ink had wider crux of helix (yellow box) and shorter intertragic notch (yellow circle) compared to those of the 3D model or the printed ear using the NC ink (Figure 3i, front view). More importantly, the overhang structure of the ear printed using the Polym ink collapsed (Figure 3i, side view) where auricular helix touched the base, whereas the ear printed with the NC ink maintained the overhang structure. After ionic crosslinking and incubation in PBS, both ears swelled to a certain extent (Figure S3,4). Still, the ear printed with the NC ink had better structural fidelity with keeping the overhang structure.

### 3.4 In vitro and in vivo biocompatibility

Biological activities of bioprinted chondrocytes were assessed. The chondrocytes were bioprinted using either the Polym ink or NC ink, and cultured *in vitro* for 28 days. After 28 days of culture, the bioprinted cells exhibited elongated morphologies, whereas at day 1 most of the cells retained round shapes in both gels (Figure 4a). The cell viability was quantified using images obtained from fluorescence staining. (Figure 4b) At all time points, no significant difference was observed between the Polym gel and NC gel. For both inks, viability right after printing was found to be ca. 80 %, which slightly decreased to ca. 70 % at day 28. Immunohistochemical staining confirmed deposition of collagens (type I and II) as well as glycosaminoglycans (GAGs, Safranin O/Fast green staining) by the bioprinted cells after 6 weeks of *in vitro* culture. (Figure 4c, left) The bioinks also supported matrix deposition by bioprinted human auricular chondrocytes in addition to bovine articular chondrocytes. (Figure 4c, right) For both cell types, no positive staining was observed for a negative IgG control (Figure S5).

**Figure 4.**
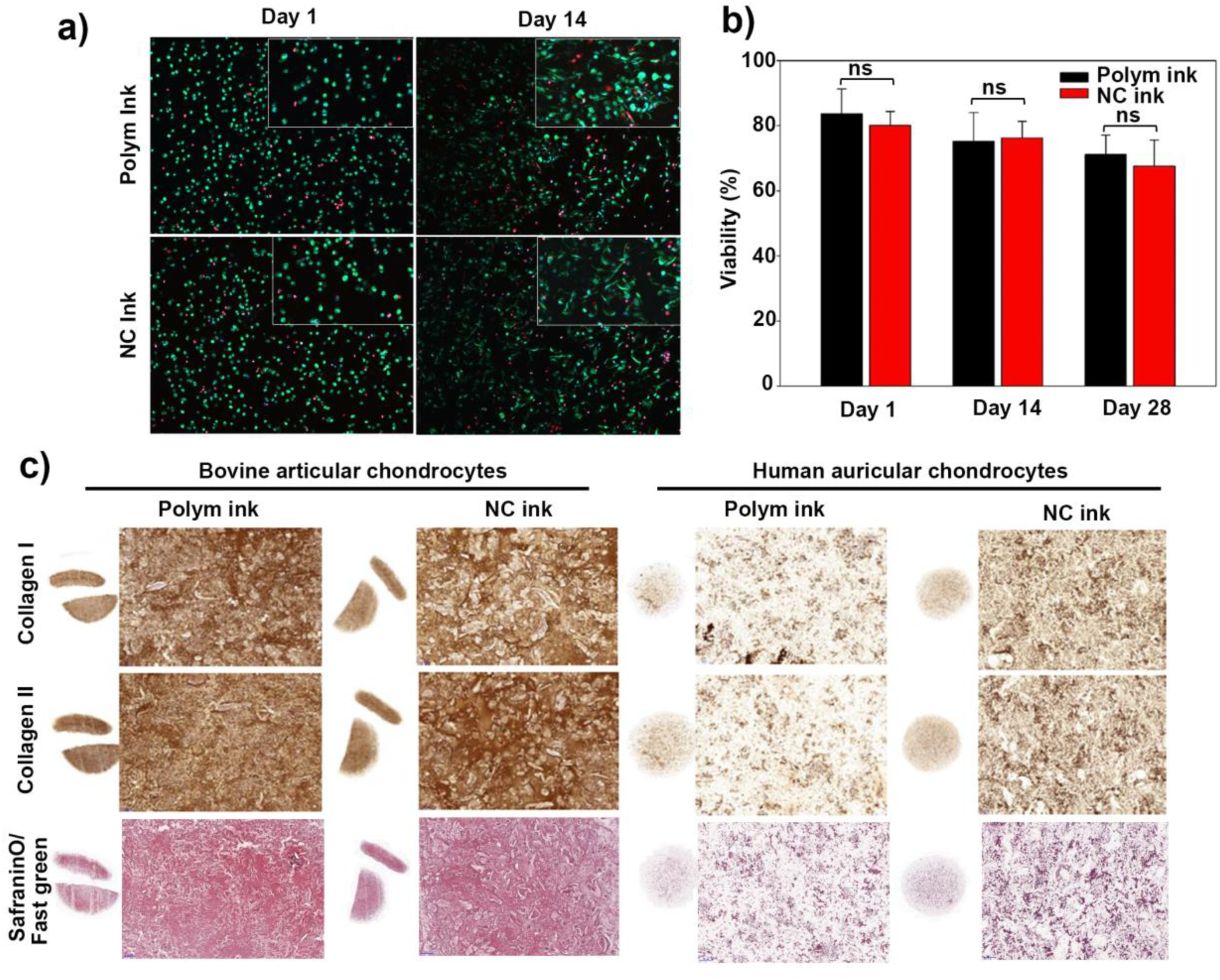
In vitro maturation of bioprinted gels. (a) Live/dead staining images of bioprinted cells. All nuclei were stained by DAPI. (b) Viability of bioprinted cells (c) Staining images of collagen I and II, and glycosaminoglycans (Safranin O staining) deposited in bioprinted gels using either bovine articular chondrocytes or human auricular chondrocytes after 6 weeks of in vitro maturation.

Finally, the bioprinted gels were implanted in subcutaneous pockets of nude mice. (Figure 5a) After 8 weeks of *in vivo* maturation, both Polym and NC gels maintained their shapes and sizes, showing that the gels are stable in a more challenging environment. (Figure 5b) Importantly, none of the gels was permissive for vessel ingrowth, as evidenced by the absence of red blood cells that display a noticeable cherry pink colour after H&E staining. For both gels, deposition of similar amounts of collagen I and II, and GAGs was confirmed, with a relatively more intense staining for collagen I compared to collagen II. (Figure 5c) In addition, calcification in the gels was not induced, which was confirmed by the negative alizarin red staining, which further highlights the biocompatibility and stability of the hydrogels.

**Figure 5.**
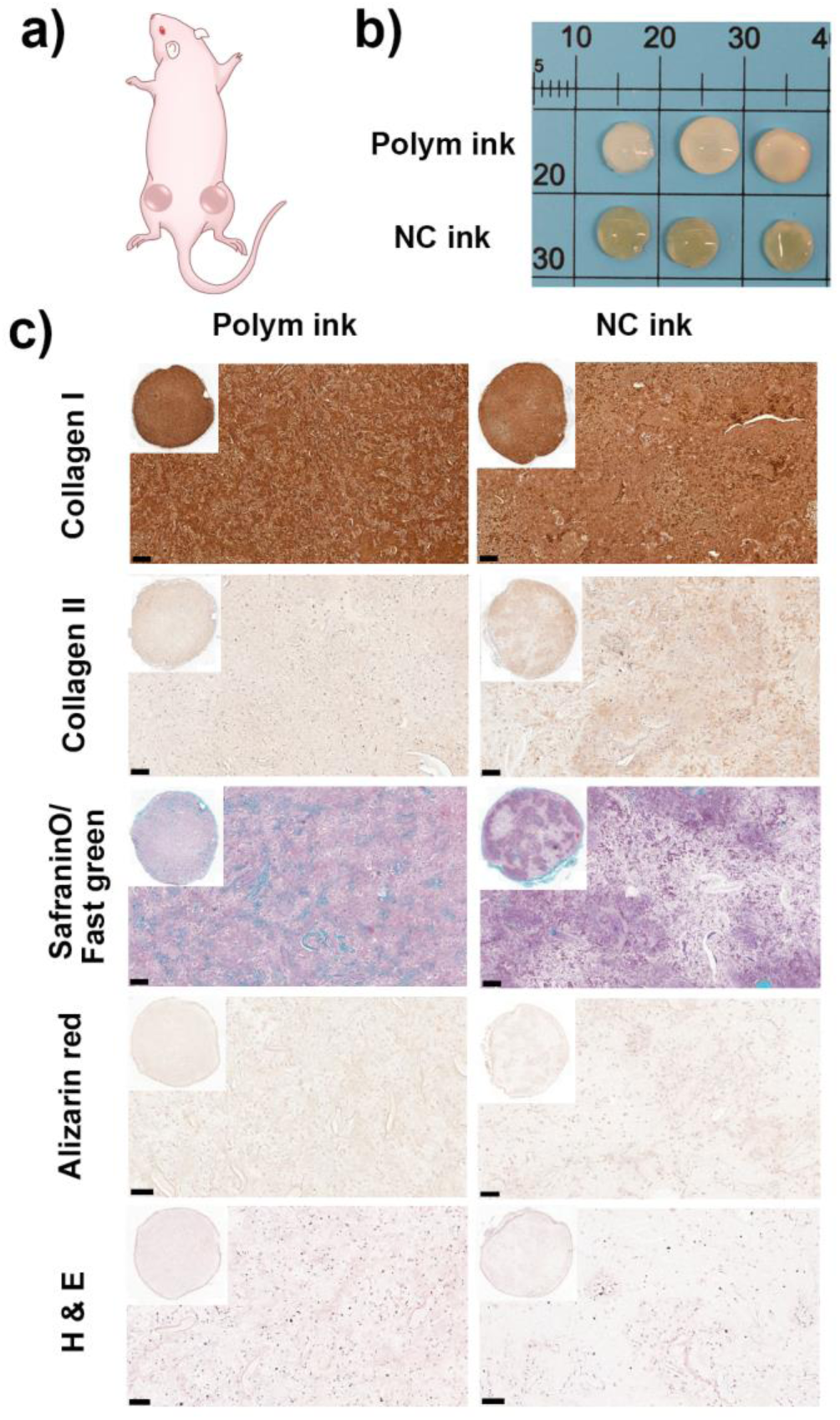
In vivo maturation of bioprinted gels. (a) Schematic presentation of the mouse subcutaneous implantation model (b) Bioprinted gels after 8 weeks of in vivo maturation (c) Immunohistochemical staining of collagen I / II and histochemical staining of glycosaminoglycans (safranin O/fast green), calcium deposition (alizarin red, no positive staining) and H & E of bioprinted gels after 8 weeks of in vivo maturation. Bar = 100 μm

## 4. Discussion

We previously demonstrated that addition of cationic SiNPs to polymeric bioinks composed of anionic polysaccharides significantly increased printability as well as printing fidelity, which was attributed to electrostatic interactions between the SiNP surfaces and polysaccharide chains.^[22]^ Although the study showed the great promise of nanocomposite-based bioinks, a high concentration of SiNPs (6 wt%) was required to achieve desirable printing properties, which could impede the use of these bioinks in clinical applications. In the present study, we aimed to develop a nanocomposite bioink by inducing dynamic covalent linkages between NPs and polymers to achieve high printing fidelity and biocompatibility using reduced amounts of NPs compared to the electrostatic interaction-based nanocomposite ink. In the NC ink we prepared here, a double network of physical crosslinking of gellan and reversible imine bonds between NH_2_-SiNPs and OxAlg mainly contribute to viscoelastic properties of the ink (Figure 1a), whereas the properties of the Polym ink are mainly controlled by gellan. The dynamic covalent bonds between the particles and polymers indeed provided the ink with increased yield stress and viscosity, and enhanced shear thinning properties with reduced amounts of nanoparticels (2 wt%) compared to electrostatic interactions. Addition of 2 wt% NH_2_-SiNPs and induction of dynamic covalent linkages led to increased yield stress from 13.8 Pa to 71 Pa (415 % increase), whereas in our previous study addition of 6 wt% NH_2_-SiNPs and induction of electrostatic interactions resulted in increased yield stress from 10.8 Pa to 51.4 Pa (376 % increase)^[22]^. Furthermore, the yield stress of the Polym ink was increased by 139 % with addition of only 0.5 wt% NH_2_-SiNPs. On the other hand, the electrostatic interaction-based nanocomposite required at least 4 wt% NPs to obtain significant rheological enhancement^[22]^. In addition, the NC ink exhibited good shear recovery (80 %), which is attributed to reversible and self-healing properties of the imine linkages. Overall, the present study demonstrated that dynamic covalent linkages between NPs and polymers in a nanocomposite could efficiently improve rheological properties desirable for quality printing.

Stabilization of a printed construct is necessary to maintain the structure of the printed construct. For the NC and Polym inks, the stabilization was induced by ionic crosslinking of alginate by Ca^2+^ ions. For both inks, compressive moduli did not decrease drastically over 7 days. However, significant decreases in tensile moduli within 6 h were observed for both inks, which should be due to diffusion of calcium ions from the gels to the buffer solution. The result implies that ionic crosslinking largely contributed to the initial tensile strength at time 0 h. Still, the presence of imine linkages enhanced both compressive and tensile strength. The NC ink not only retained higher moduli at 0 h but better stability for 7 days compared to the Polym ink. The enhanced mechanical stability is attributed to the relatively stable nature of covalent imine linkages compared to Ca^2+^-alginate ionic bonds.^[38]^

We showed that enhanced rheological properties led to improved printing precision and fidelity. Yield stress has been reported as an indicator for good printability, where materials with higher yield stress retain better printing fidelity with a less collapse of overhang structures and fusion between neighboring filaments compared to the ones with lower yield stress.^[39]^ Indeed, the NC ink, which retained higher yield stress than the Polym ink, allowed for printing porous structures with less collapses than the Polym ink. In addition, both increased yield stress and viscosity by NH_2_-SiNPs led to significantly suppressed diffusion of the ink at intersections in a grid, resulting in high printing fidelity. With enhanced printability and printing fidelity, a centimetre-scale multi-layered grid structure as well as an ear structure with overhangs were successfully printed using the NC ink.

The NC ink supported spreading and matrix deposition of bioprinted chondrocytes as efficiently as the Polym ink did. Cartilage is one of the most studied tissue in bioprinting because it contains only single cell type and is not vascularized so that the tissue is considered to be relatively easy to fabricate compared to other tissue.^[40]^ In addition, there are high clinical demands for generation of patient-tailored articular or auricular cartilage to treat a number of cartilage-related diseases such as microtia/anotia, osteoarthritis or traumatic cartilage injuries.^[41, 42]^ However, *in vitro* promotion of matrix deposition by chondrocytes isolated from native tissue is often challenging in many hydrogel or composite systems.^[41]^ Furthermore, for a bioink, the mechanical properties of the material should be suitable not only to support growth of chondrocytes but also for providing high printability. In the present study, we showed that the addition of NPs in the ink did not compromise the biocompatibility of the Polym ink, where intense staining for collagens and GAGs was observed after 6 weeks of *in vitro* culture of bioprinted bovine articular chondrocytes. It is known that increased crosslinking density could impede chondrogenesis and/or induce hypertrophy and calcification due to limited diffusion of nutrients/waste as well as restricted matrix remodelling by the cells.^[43, 44]^ However, in the printed NC gels, we observed deposition of large amounts of matrix which was uniformly distributed throughout each gel. In addition, there was no significant difference in intensity or uniformity of staining between the Polym and NC gels despite of higher mechanical strength of the NC gel than that of the Polym gel. This observation is attributed to reversible nature of imine linkages which allows for remodelling of matrix via breakage and re-formation of the bonds. ^[38]^ Bioprinted human auricular chondrocytes, however, did not produce as much matrix as bovine articular chondrocytes, which could be mainly attributed to lower cell density (5*10^6^ cells/mL) in the gels compared to bovine cells (10*10^6^ cells/mL) due to limited amounts of clinical samples. It is known that the efficiency of chondrogenesis in a hydrogel increases with increasing cell density in the gel.^[45, 46]^ In the study by Sun et al, for example, ~9 times higher amounts of GAG deposition was observed for the cell density of 20 * 10^6^ cells/mL compared to that of 8 * 10^6^ cells/mL in a hydrogel composed of PLLA and PEG after 8 weeks of *in vitro* culture. Therefore we consider that the chondrogenesis by the auricular chondrocytes in the NC gels could be further optimized via increasing cell density.

## 5. Conclusion

We demonstrated that formation of dynamic covalent linkages between NPs and polymers in a nanocomposite can provide enhanced mechanical properties and printing fidelity without compromising biocompatility of the polymeric ink. Imine linkages in the NC ink led to enhanced shear thinning properties as well as increased yield stress, resulting in 3D printing of various structures with high fidelity with reduced diffusion or collapses of ink materials. Furthermore, the NC ink supported both *in vitro* and *in vivo* maturation of bioprinted chondrocytes, where intense matrix deposition was confirmed with good mechanical stability of the bioprinted gels. Considering the availability of aldehyde presenting biopolymers including many oxidized polysaccharides, we expect the strategy could be expanded to other polymeric bioinks to provide desirable printing and mechanical properties, leading to practical applications of bioprinted tissue.

## Acknowledgements

This work was financially supported by Sinergia grant from the Swiss National Science Foundation (CRSII5_173868 to MZW). The authors acknowledge the use of the Scientific Center for Optical and Electron Microscopy facilities of ETH Zürich, the support by Dr. Martin Badescher for ATR-FTIR measurements and by Mr. David Fercher for isolating human auricular chondrocytes.

