## Supplemental data for "Nanocomposite bioink exploits dynamic covalent bonds between nanoparticles and polysaccharides for precision bioprinting"

**Supplementary Information**

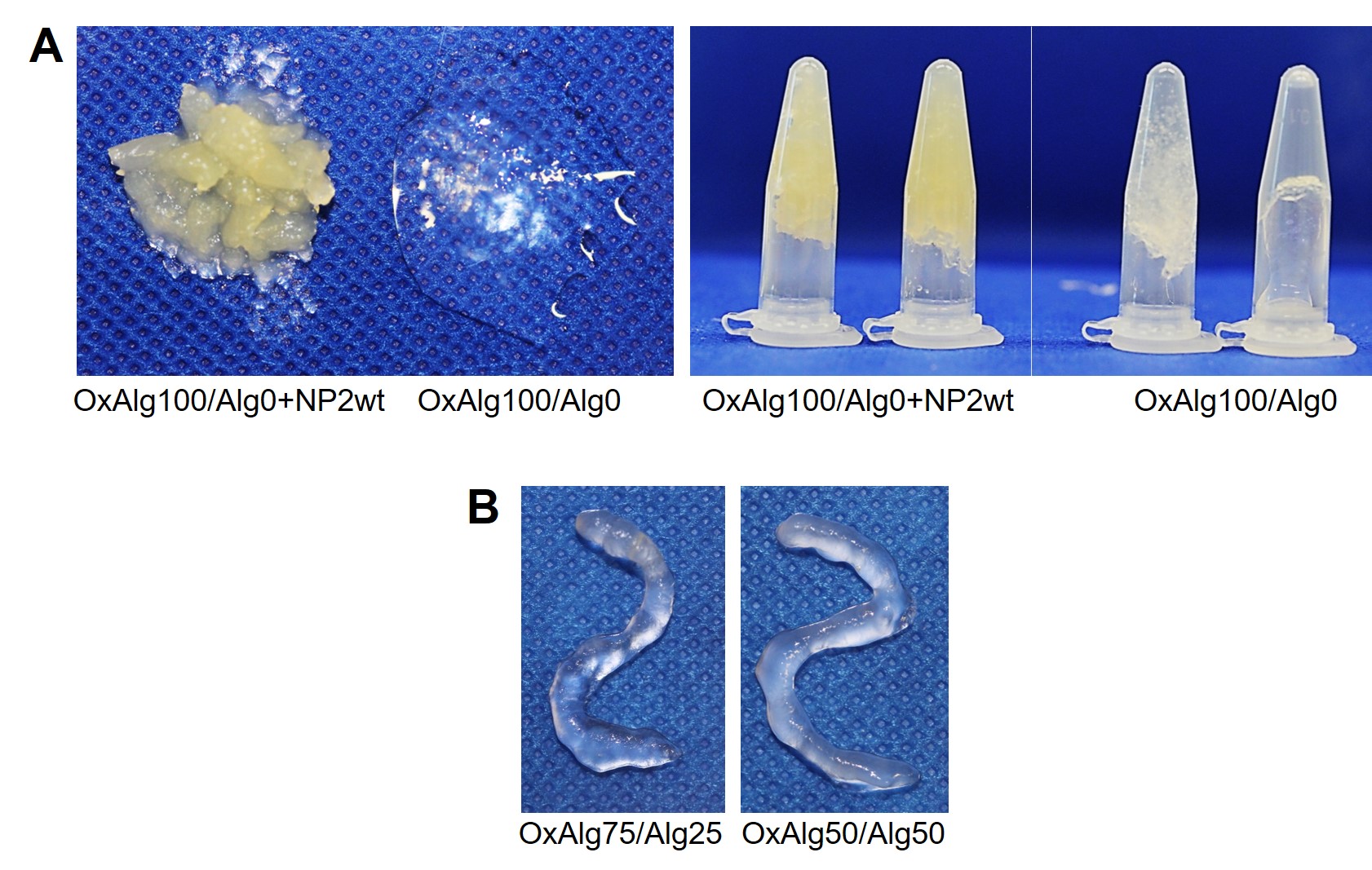

**Figure S1**. Photographs of inks with vaying ratios of OxAlg to alginate (Alg). A) The inks with the ratio of 100:0 (OxA:Alg; 3%/0%) with/without nanoparticles. B) The inks with the ratio of 75:25 (2.25%/0.75%) and 50:50 (1.5%/1.5%) without nanoparticles.

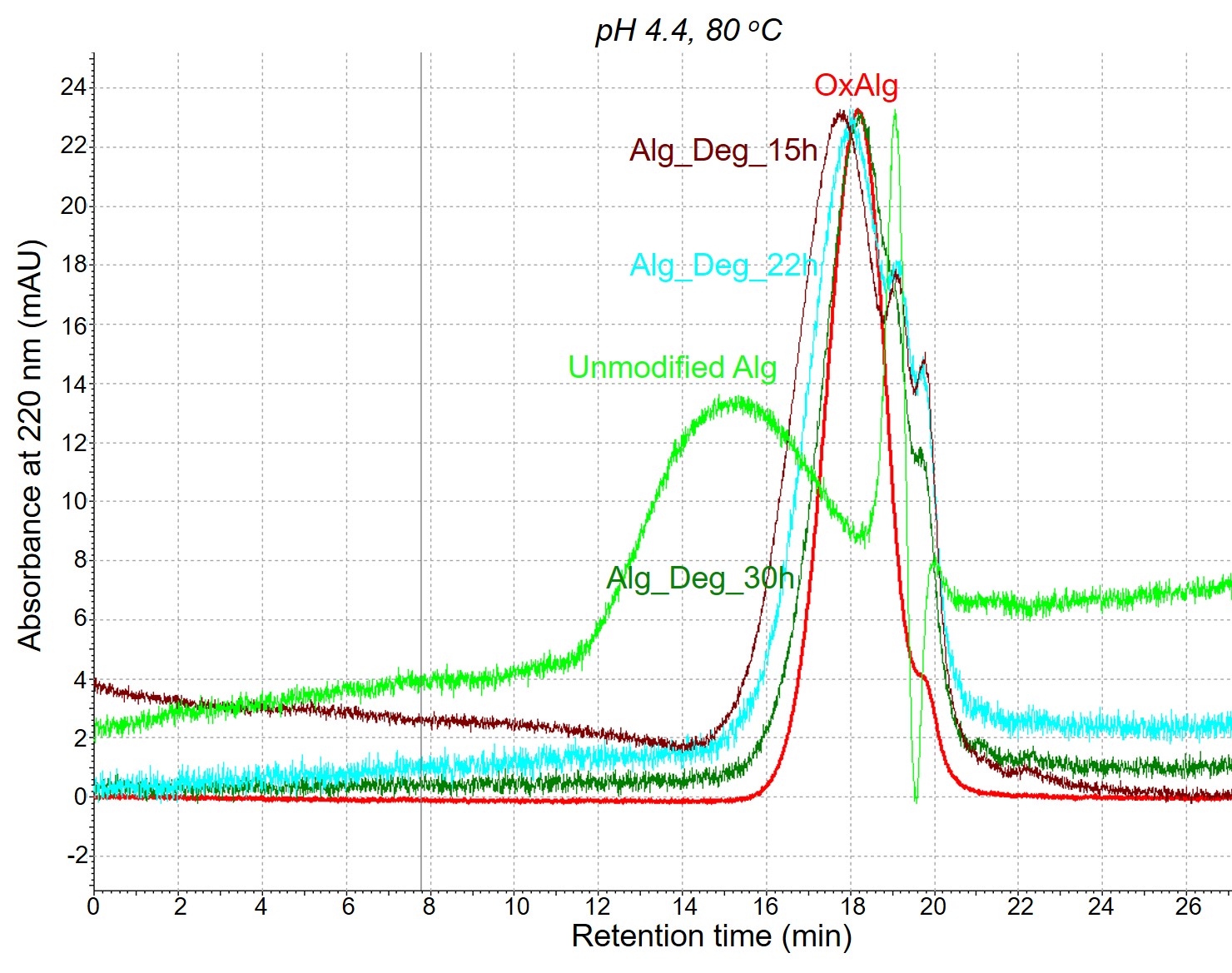

**Figure S2**. GPC spectra of alginate, OxAlg and depolymerized alginate (Alg-Deg) (15, 22, 30 h)

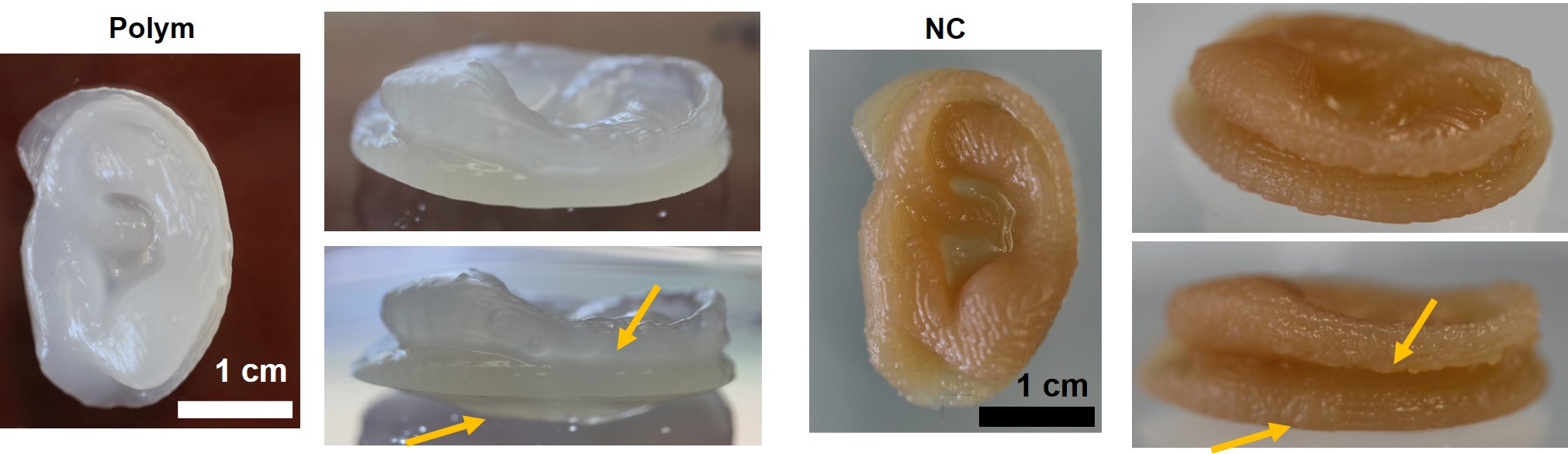

**Figure S3**. Photographs of printed ears after crosslinking overnight. Arrows indicate the regions where differences in structural fidelity were observed prominently between two ears printed using Polym or NC ink.

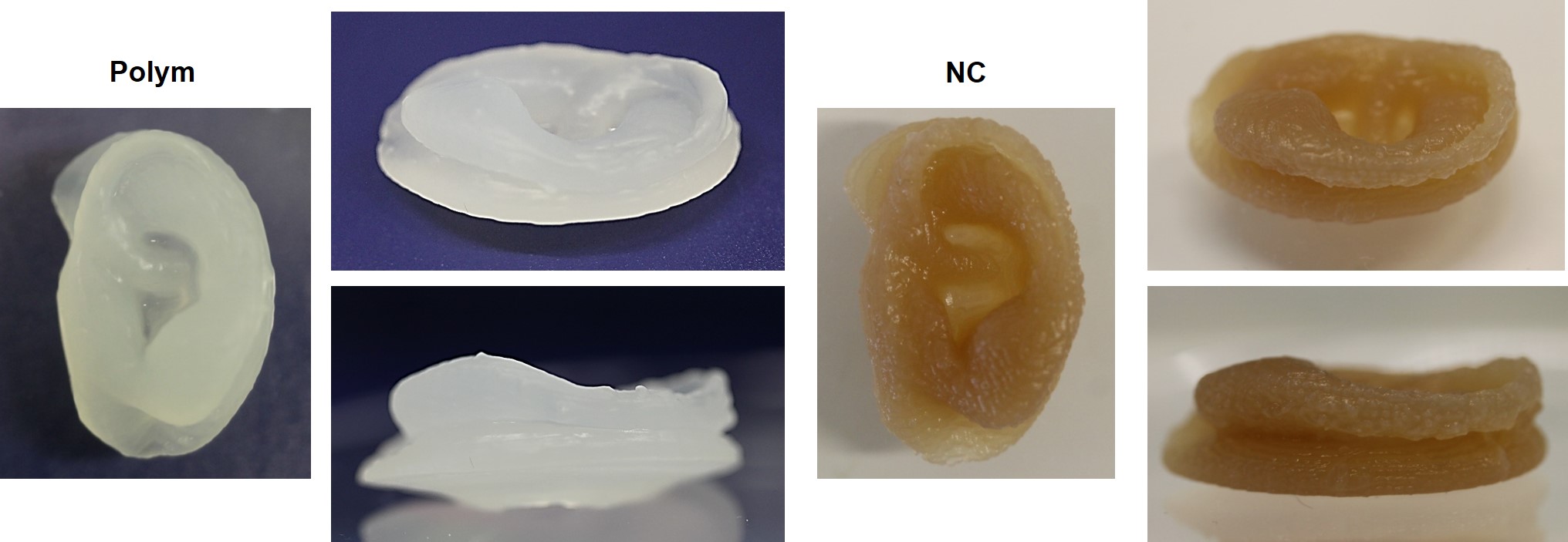

**Figure S4**. Photographs of printed ears after swelling in PBS for 72h

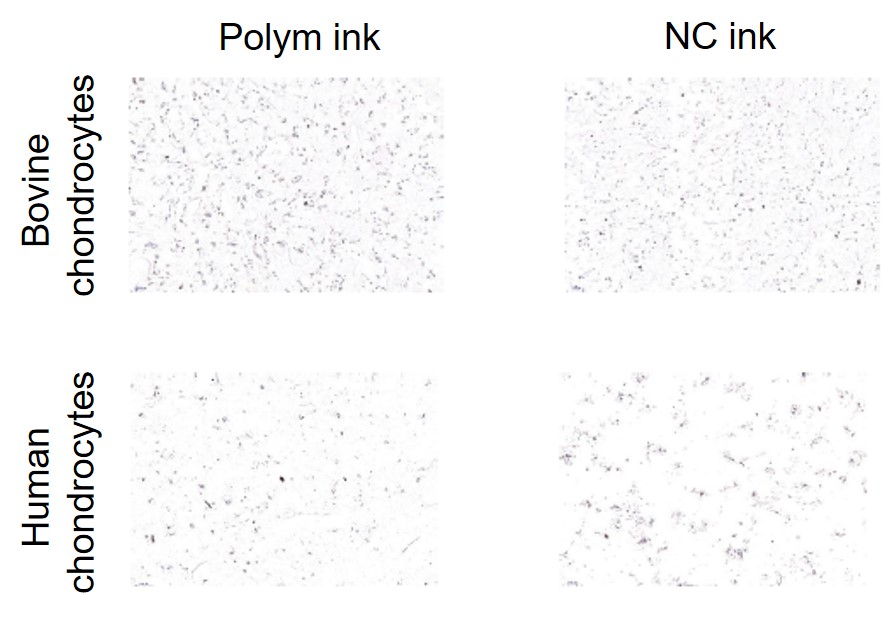

**Figure S5**. IgG staining of the gels as a negative contol
